# Single-Cell RNA-Seq Reveals Novel Mitochondria-related Musculoskeletal Cell Populations during Adult Axolotl Limb Regeneration Process

**DOI:** 10.1101/704841

**Authors:** Tian Qin, Chun-mei Fan, Ting-zhang Wang, Long Yang, Wei-liang Shen, Heng Sun, Jun-xin Lin, Magali Cucchiarini, Nicholas D. Clement, Christopher E. Mason, Varitsara Bunpetch, Norimasa Nakamura, Rameah Bhonde, Nicholas D. Clement, Zi Yin, Xiao Chen

**Affiliations:** Dr. Li Dak Sum-Yip Yio Chin Center for Stem Cells and Regenerative Medicine and Department of Orthopedic Surgery of The Second Affiliated Hospital, Zhejiang University School of Medicine, Hangzhou, China; Dr. Li Dak Sum-Yip Yio Chin Center for Stem Cells and Regenerative Medicine, Zhejiang University School of Medicine, Hangzhou, China; Key Laboratory of Tissue Engineering and Regenerative Medicine of Zhejiang Province, Zhejiang University School of Medicine, Hangzhou, China; Zhejiang University-University of Edinburgh Institute & School of Basic Medicine, Zhejiang University School of Medicine, Hangzhou, China; Department of Sports Medicine, Zhejiang university School of Medicine, Hangzhou, China; China Orthopedic Regenerative Medicine Group (CORMed), Hangzhou, China; Key Laboratory of Microbial Technology and Bioinformatics of Zhejiang Province, Hangzhou, China.; Department of Orthopedic Surgery of the Second Affiliated Hospital, Zhejiang University School of Medicine, Hangzhou, China; Center of Experimental Orthopaedics, Saarland University Medical Center and Saarland University, Kirrbergerstr. Bldg 37, D-66421 Homburg/Saar, Germany; Department of Orthopaedics and Trauma, Royal Infirmary of Edinburgh, Little France, University of Edinburgh, Edinburgh, Scotland, UK; Department of Physiology and Biophysics, Weill Cornell Medicine, New York, NY, USA; The HRH Prince Alwaleed Bin Talal Bin Abdulaziz Alsaud Institute for Computational Biomedicine, Weill Cornell Medicine, New York, NY, USA; The WorldQuant Initiative for Quantitative Prediction, Weill Cornell Medicine, New York, NY, USA; The Feil Family Brain and Mind Research Institute, Weill Cornell Medicine, New York, NY, USA; Institute for Medical Science in Sports, Osaka Health Science University, 1-9-27, Tenma, Kita-ku, Osaka city, Osaka 5300043, Japan; Dr.Ramesh Bhonde, Director Research, Dr. D. Y. Patil Vidyapeeth, Pimpri, Pune 411018, India

## Abstract

While the capacity to regenerate tissues or limbs is limited in mammals including humans, unlike us, axolotls are able to regrow entire limbs and major organs. The wound blastema have been extensively studied in limb regeneration. However, due to the inadequate characterization and coordination of cell subpopulations involved in the regeneration process, it hinders the discovery of the key clue for human limb regeneration. In this study, we applied unbiased large-scale single-cell RNA sequencing to classify cells throughout the adult axolotl limb regeneration process. We computationally identified 7 clusters in regenerating limbs, including the novel regeneration-specific mitochondria-related cluster supporting regeneration through energy providing and the COL2+ cluster contributing to regeneration through cell-cell interactions signals. We also discovered the dedifferentiation and re-differentiation of the COL1+/COL2+ cellular subpopulation and uncovered a COL2-mitochondria sub-cluster supporting the musculoskeletal system regeneration. On the basis of these findings, we reconstructed the dynamic single-cell transcriptome atlas of adult axolotl limb regenerative process, and identified the novel regenerative mitochondria-related musculoskeletal populations, which yielded deeper insights into the crucial interactions between cell clusters within the regenerative microenvironment.

## INTRODUCTION

The ability to fully regenerate damaged and lost tissues would be of tremendous benefit for clinical medicine; however, the limited regeneration capacity of many human tissues, including limbs, presents a formidable clinical hurdle *(Shieh and Cheng, 2015)*. The axolotl is a widely used model organism in limb regeneration due to its powerful ability to reconstitute a fully functional limb after amputation throughout adulthood. After amputation, a blastema forms which is defined as a mass of progenitor cells that is capable of regenerating of limbs by autonomously patterning itself to precisely replace the lost structures *(Brockes, 1997)*. Histological analysis and lineage tracing studies illustrated that blastema cells originate from the mesodermal tissues including dermal fibroblasts, Schwann cells, and myogenic cells *(Flowers et al., 2017)*. Meanwhile, many studies based on transcriptomes, gene editing and proteomic analysis provided much information concerning the key molecular mechanisms in blastema formation *(Looso et al., 2013; Nacu et al., 2016; Haas and Whited, 2017)*. The oncogenes were reported to burst in the early stage *(Stewart et al., 2013)* and two crucial genes *cirbp* and *kazald1* were discovered upregulated in blastemas *(Bryant et al., 2017)*. MARCKS-like protein was reported to be an initiating molecule *(Sugiura et al., 2016)* and some proteins such as CCDC88c and DIXDC1, were discovered to be involved in Wnt signaling during blastema formation *(Rao et al., 2009)*. However, despite this accumulated knowledge, the following critical aspects of the axolotl regenerative mechanism remain unclear: cell populations and their interactions that contribute to blastema constitution, and whether the cartilaginous skeleton is related to musculoskeletal system regeneration. Furthermore, the differences between axolotl and human cell populations and their related functions are also not well understood. Thus, a more precise and detailed cell population analysis during blastema development is of critical importance.

While prior work has shown that the blastema is a heterogeneous pool of progenitor cells, reflecting different propensities for lineage differentiation, further identification of different cell populations that contribute to limb regeneration is needed. Traditional sequencing methods using whole tissues, mask potential cell heterogeneity within the tissue and may add bias to be understanding of the cellular hierarchy. Single-cell RNA-sequencing (scRNA-seq) analysis allows unbiased and high-throughput analysis of gene expression profiles from an individual cell level in order to investigate cell population heterogeneity *(Lanctôt, 2015)*. ScRNA-seq reveals novel cell types and reconstructs lineage hierarchies of tissue types such as human pluripotent stem cells *(Han et al., 2018a)*, blood dendritic cells *(Villani et al., 2017)*, neurons *(Li et al., 2017)*, and uterus epithelium *(Wu et al., 2017)*, whereafter it can even maps an atlas *(Han et al., 2018b)*. Recently, the scRNA-seq was performed on axolotl to uncover the connective tissue subpopulations with their characters and mesenchymal cellular diversity during regeneration *(Gerber et al., 2018; Leigh et al., 2018)*. However, the alterations of global limb cell populations during regeneration process were not clarified.

In this study, scRNA-seq was performed on blastema cells collected limb tissues at four stages during the post-amputation period of adult axolotls, from regeneration early to late stages. The transcriptome profiling displayed a comprehensive axolotl regeneration cell atlas by clustering cells across the regeneration time course and reconstructing connectivity map between each cell cluster. We also reconstructed the regeneration pseudo-temporal ordering and found the dynamics of the musculoskeletal system cells during the regeneration process. We also identified a novel mitochondria-related cell population which plays a vital energy supporting character in regeneration. All of these provide a greater understanding of the limb regeneration process.

## RESULTS

### Isolation of blastema cells after amputation and pre-processing of sequencing data

Samples were isolated from 1 mm *(Voss et al., 2015)* of adult axolotl forelimbs (0 dpa) and from 1 mm of heterogeneous tissue collected from the regenerated distal limb tips at 3 days, 7 days, and 21 days post amputation (3 dpa, 7 dpa, 21 dpa) which represent the early, middle, and late stage of regeneration respectively (Fig. 1A, upper left; fig. S1A). These samples were prepared as single-cell samples for sequencing using Fluidigm C1 system with HT IFCs (Fig. 1A, upper middle and right) which could detect an average of 5000 genes per cell.

**Figure 1.**
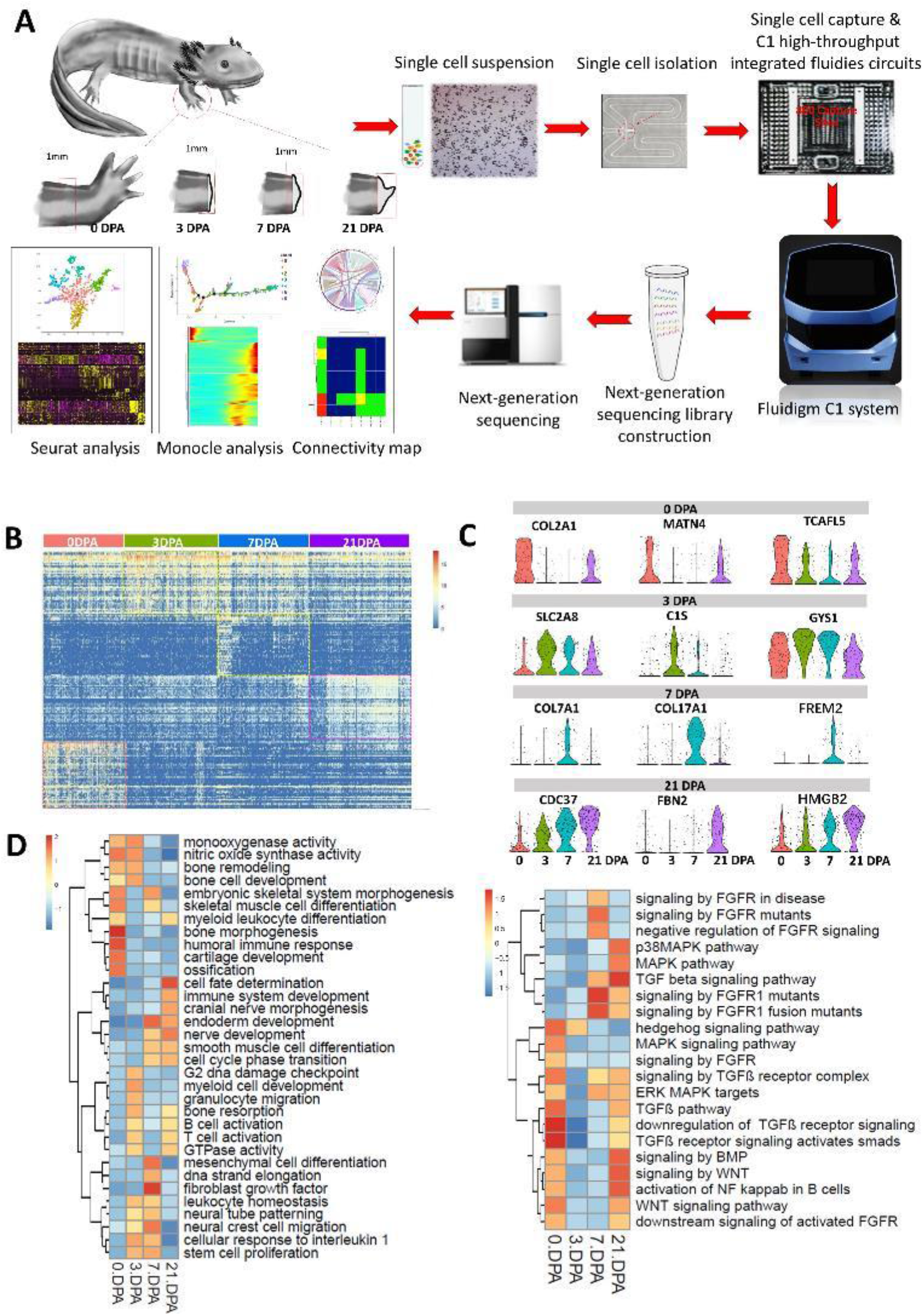
Single-Cell RNA-Seq Reveals changes along axolotl limb regeneration the time courses. (A) Schematic diagram of the blastemal time course experiment. The red box on limbs indicated the amputation location on axolotls’ limbs (upper left). The pictures shows that the amputated tissues were trypsinized to yield a single-cell suspension (upper middle) and trapped using the Fluidigm C1 auto prepare system (C1 high-throughput integrated fluidics circuits (HT IFCs) and HiSeq systems) for the downstream single-cell mRNA-seq experiment. (B) Heatmap of 936 single cells from the four regeneration stages by transcriptome analysis. DPA, day post-amputation. (C) Violin plots of marker genes from 4 time points. (D) Heatmaps of GO terms (left) and signaling pathways (right) by GSEA analysis of each time point post-amputation upregulated genes. Bars of heatmaps are color-coded according to p-value, ranging from low value (blue) to the highest value (red), according to the adjacent color key.

Our sequencing data were mapped with three *de novo* transcriptome assemblies from previous studies containing both axolotl and Chinese giant salamander, in addition to the genome of *Xenopus laevis (Stewart et al., 2013; Bryant et al., 2017; Geng et al., 2017)*. From these alignments, we received 609,180 transcripts. The functional annotation of de novo assembled transcriptomes was performed by Trinotate (http://trinotate.github.io). After data quality control and removal of 72 multiple cell samples, 936 cells with >212,000 transcripts (15650 genes) were selected for further analysis (fig. S1 B and C). Recently, Nowoshilow et al. reported the sequencing and assembly of the axolotl genome *(Nowoshilow et al., 2018)*, which has been very valuable traditional RNA sequencing researches. We mapped our data sets to the genome with a 90% mapping rate, confirming the high reliability of our RNA library (fig. S1D). However, since our sequencing used the mainstream 3’ end enrichment library method and the published genome and transcriptome have incomplete gene annotation of the 3’ end sequence, the published data is unsuitable for our high-throughput single-cell sequencing. Full-length sequencing is required to establish a full-length transcriptome to match the genome.

### Dynamic patterns of gene expression along axolotl limbs regeneration time course

Evidence provided by previous studies demonstrated dramatic changes in the transcriptome at different time points during post-amputation *(Stewart et al., 2013; Wu et al., 2013)*. In order to understand the transcriptome changes during regeneration, we analyzed differentially expressed (DE) genes at each time points after amputation and our results also showed discernible differences in gene expressions of the four stages (Fig.1B; Fig. S1E; Table S1). Cartilage associated genes *COL2A1*, *MATN4* and *TCAFL5* (Fig. 1C) were highly expressed in cells at 0 dpa, while glucose metabolism genes SLC2A8, C1S, and GYS1 were highly expressed in cells at 3 dpa. At 7 dpa, epidermis associated genes *COL7A1*, *COL17A1*, and *FREM2*, and an upper limb development key gene *TBX5* (T-box 5) *(Simon et al., 1997)* were highly expressed. Lastly at 21 dpa, *CDC37*, a cell division cycle protein gene, *FBN2*, a component of connective tissue microfibril gene, and *HMGB2*, a cell death programmed gene, which are also characterized by the high cell proliferation and multi-lineage development ability were highly expressed (Fig. 1C).

To further obtain a functional insight of the cells in each time point, we performed gene set enrichment analysis (GSEA); According to the GSEA results, cells from each time point possessed unique gene ontology (GO) and key signaling pathways (Fig. 1D, fig. S1F). Results revealed that GO terms of 0 dpa cells were related to cartilage development and morphogenesis while 3 dpa cells were related to functions involving cell proliferation, monooxygenase activity, bone remodeling, and cellular response to interleukin 1. In the later stage on 21 dpa, the blastema displayed GO terms associated with endoderm development, mesenchymal differentiation, limb morphogenesis, cell fate determination, and fibroblast growth factor (Fig. 1D, left). For the key signaling pathways, GSEA results illustrated that the MAPK and FGFR pathway mainly participated during the middle and late stages of regeneration, while TGFb, Wnt, and BMP pathways were activated at a more mature stage (21 dpa and unamputated limbs) (Fig. 1D, right). These results demonstrate the integral dynamic patterns of gene expression along the axolotl limb during according regeneration time.

### Cell heterogeneity in blastemas of different amputation time points by single-cell profiling

General differentiated expression analysis probably masks the cell heterogeneity within each time point. To identify cell types of axolotl limb regeneration, we used Seurat to perform principal component analysis (PCA) and t-distributed stochastic neighbor embedding (t-SNE) analysis *(Kiselev et al., 2017)*. Unsupervised clustering based on principal components of the most variably expressed genes was used to separate cells (4 time points) into 7 clusters, which were visualized using t-SNE (Fig. 2A, Fig. S2A) with each cluster possessing a unique set of signature genes (Fig. 2B, Table S2; Fig. 2, D-J, left). Each group has a different cell quantity proportion at different time points (Fig. 2C, Fig. S2B). GO and signaling pathways analysis were performed on upregulated genes of each cluster using GOEAST (http://software.broadinstitute.org/gsea/index.jsp) (Fig. 2, D-J, right, Fig. S2C). Based on the marker genes and GO analysis results, the 7 clusters (C1-C7) were labeled as the general cluster, mitochondria cluster, limb morphogenesis cluster, chondrocyte cluster, inflammation cluster, cell proliferation cluster, and apical epithelium cap (AEC) cluster.

**Figure 2.**
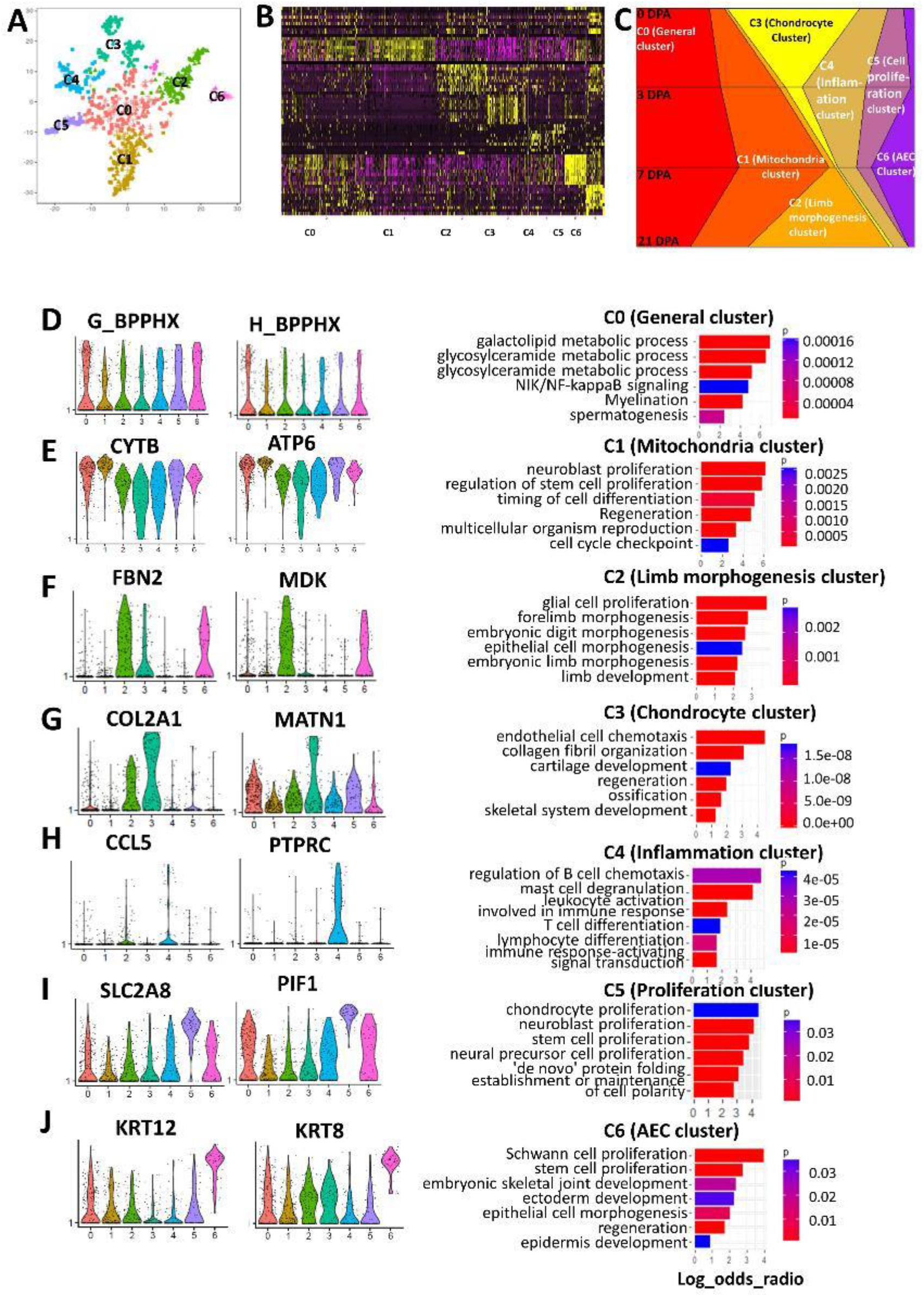
Single-Cell Profiling Reveals Cell Heterogeneity in axolotl limb tissues from different regeneration stage. (A) T-distributed stochastic neighbor embedding (t-SNE) visualizations of all cell clusters identified using the computational pipeline (B) Heatmap of top 20 differentiated gene of 7 clusters. (C) The proportion of cells at each time point in each cluster. (D-J) Violin plots of marker genes and GO analysis from 7 cluster (left). Enrichment log odds ratio for each GO term are shown as histograms (right). Bars color-coded according to p value, ranging from low detected to the highest detected levels (blue and red).

#### C0 (General cluster)

This cluster was generally located at the center of all clusters and was similarly expressed in all four dpa cells. Marker gene functions were consistent and GO terms were dispersed (Fig. 2D, Table S3).

#### C1 (Mitochondria cluster)

The mitochondria cluster was shown in high proportion in cells from regenerating stages (3 dpa, 7 dpa and 21 dpa) when compared to cells from 0 dpa (Fig. 2C, Table S4). Differential analysis demonstrated that almost all the top differential genes were mitochondria-related genes including *CYTP* and *ATP6* (Fig. 2E, left). In order to gain more knowledge on this particular cluster, we removed mitochondrial genes and re-evaluated the differences between the mitochondrial cluster and other clusters, because mitochondrial genes were significantly expressed, possibly masking other highly expressed genes. After removing mitochondria-associated genes, GO terms of the remaining upregulated genes were mainly related to regeneration and development of neuron and stem cells as well as regulation of programmed cell death (Fig. 2E, right). These results are consistent with recent studies demonstrating that mitochondrial activity influences the cell differentiation *(Schieke et al., 2008)* and response to stress (Garrett-Bakelman et al., 2019).

#### C2 (Limb morphogenesis cluster)

This cluster highly expressed *FBN2* (Fibrillin 1) and *MDK* (Midkine) with the main function involving forelimb morphogenesis such as embryonic limb morphogenesis, epithelial cell morphogenesis, and embryonic digit morphogenesis (Fig. 2F, Table S5).

#### C3 (Chondrocyte cluster)

Cells in this cluster were mainly isolated from 0 dpa limbs and highly expressed cartilage-related genes, including *COL2A1* and *MATN1* (Fig. 2G, left); GO term results were prominently related to skeletal and cartilage development (Fig. 2G, Fig. S2 C, Table S6). This cluster represented the cartilaginous skeleton in unamputated limbs, and cartilage was reported as the source of regenerative skeleton in lizards and frog *(Lozito and Tuan, 2016, 2017)*. This indicats that the chondrocyte cluster may play a role in limb skeleton regeneration.

#### C4 (Inflammation cluster)

These cells were largely involved in immune response, including leukocyte activation, and B cell and T cell activation with the marker genes *PTPRC* (protein tyrosine phosphatase, receptor type C) and *CCL5* (Fig. 2H, Table S7). This cluster represented the inflammation cells that exist in axolotl limbs and are responsible for providing a cohort of important growth factors and signaling molecules after injury, which not only regulate new vessel growth but also modulate scarring responses *(Godwin et al., 2013)*.

#### C5 (Cells proliferation cluster)

GO analysis results showed that this cluster displayed a strong proliferation characteristic with marker genes *SLC2A8* and *PIF1*, in addition to well as the terms of stem cell proliferation, neuroblast proliferation, and chondrocyte proliferation (Fig. 2I, Table S8). A previous study showed that chondrocytes near the amputation plane undergo division and proliferation responding to injury, and that these responses are limited in the late stage of regeneration. This cluster may represent a group of cells that rapidly divides and proliferates in the early stages of regeneration after amputation *(Currie et al., 2016)*.

#### C6 (AEC cluster)

Immediately after amputation, a wound epithelium (WE) was formed by migrating epithelial cells *(Roensch et al., 2013)*; it would interact with the stump tissues to form an apical epithelium cap (AEC) which is essential in creating a regenerative environment for blastema *(Christensen and Tassava, 2000; Endo et al., 2004)*.This cluster of cells highly expressed epidermis marker genes *KRT12* and *KRT8* (Fig. 2J, left), and blastema marker genes such as *PRRX1* and *CRIBP* (fig. S2D) which represent the AEC formed by the migrating epithelial cells. However, the GO term revealed that this cluster was related to not only epidermis development but also Schwann cell and stem cell proliferation (Fig. 2J, right, Table S9). These results demonstrated the traits of the AEC cluster which include inducing the differentiation in the underlying stump tissue, attracting cells accumulating below and being in response to nerve signals *(SINGER and INOUE, 1964)*.

These results provide a comprehensive classification and definition of total cells from unamputated limbs and regenerated blastemas.

### The mitochondria cluster supported cell division in regenerative limbs

To confirm the mitochondria activity in regeneration process, we detected the mitochondrion-marker TOMM20 using immunochemistry staining. The TOMM20 had the highest expression in blastema of the 3 dpa tissues. We also found that TOMM20 had higher expression level in the skin and muscle of the 3 dpa tissues compared to the 0 dpa tissues (Fig. 3A). Additionally, TEM results of epidermis (fig. S3A) indicated that the average number of mitochondria per cell in regenerative tissues were significantly higher than in normal limb tissues (Fig. 3B). These results indicated that the higher mitochondria activities in mitochondria cluster play a key role in regeneration process. To recognize the functions of mitochondria cluster, GSEA analysis was performed on it and the rest cluster. The results showed citrate cycle TCA cycle pathway were upregulated in mitochondria cluster, which point at the reliability of the identity of mitochondria cluster. The cell division-associated signaling pathways including signaling by ROBO receptor, G alpha1213 signaling events, apoptotic execution phase, the energy metabolism-associated signalings such as signaling by RHO GTPases, and cell death regulation-associated pathways including Nrage signals death through JNK were upregulated in mitochondria cluster (Fig. 3C). These indicated that the mitochondria cluster supported cell division and regulated cell death in regenerative tissues.

**Figure 3.**
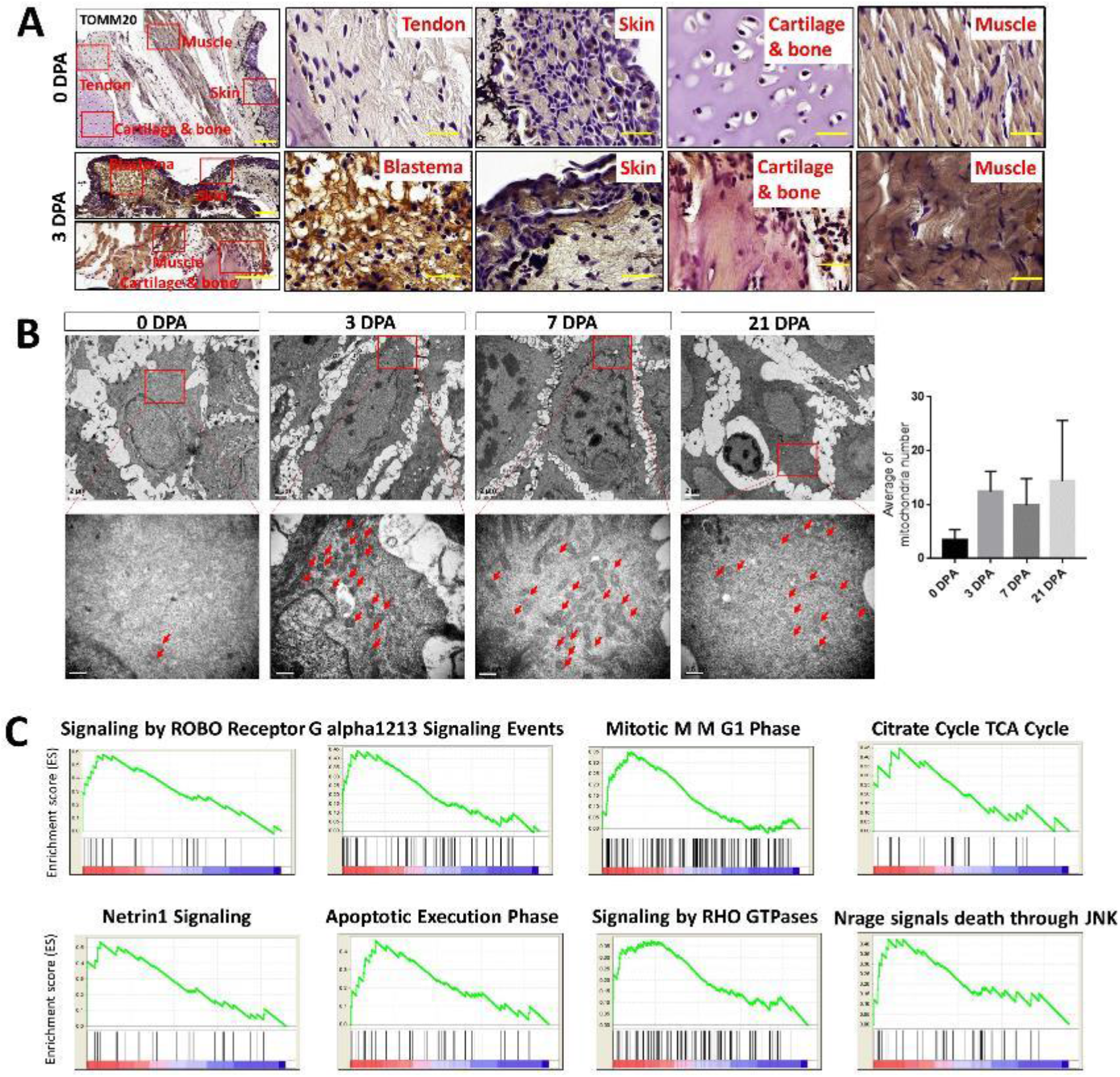
The mitochondria cluster specifically existed in regenerative limbs. (A) Immunostaining of TOMM20 in tendon, skin, cartilage and muscle of 0 dpa tissues, and blastema, skin, cartilage and muscle of 3 dpa tissues. Scale bar, 50 μm. (B) TEM of regenerative samples at each time points. Scale bar, upper 2 μm, lower 0.5 μm. (C) The top 8 signaling pathways in mitochondria cluster compared with the rest clusters by GSEA signaling pathway analysis.

### The microenvironment regulating regeneration process of different cell clusters by connectivity map

Different cell clusters co-exist within the same environment and physically surround each other and communications between these cells clusters may also play a part in regulating cell state and determining cell fate. Thus, we next constructed a connectivity map of the 7 sub-clusters for each corresponding time point respectively in order to reveal the potential interaction of different clusters by using known ligand-receptor pairs *(Camp et al., 2017; Zepp et al., 2017)*. We detected 49 connections of 928 ligand-receptor pairs from 7 sub-clusters in 0 dpa, 3 dpa and 7 dpa cells and 36 connections of 928 ligand-receptor pairs from 6 sub-clusters in 21 dpa (Fig. 4A; fig. S3A).

**Figure 4.**
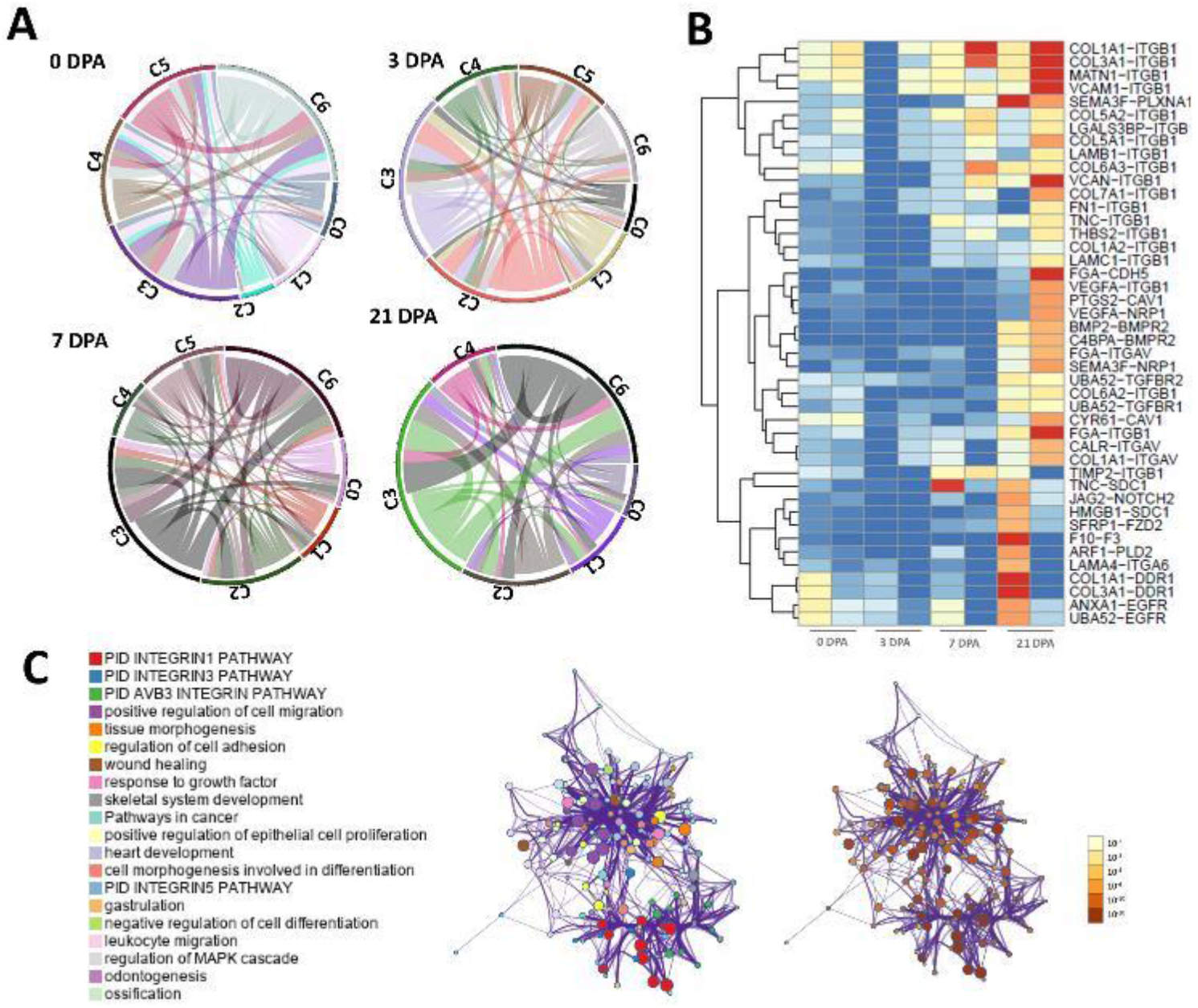
Connectivity maps of 7 clusters during the time course. (A) Circle plot depicts the total number of ligand-receptor interactions between each sub-cluster in each time point post amputation. (B) Top receptor-ligand expression between chondrocyte cluster and AEC cluster. (C) GO terms of top receptor-ligand genes between chondrocyte cluster and AEC cluster.

The connectivity map revealed that the C3 had the most ligands to C6 when compared to other cell populations in all 4 temporal groups (Fig. 4A, Fig. S3B), thus showing a potential contribution of chondrocytes to AEC development (Fig. 4A; fig. S3B). Further analysis of the ligand-receptor pair interactions between C3 and C6 showed abundant extracellular matrix (*COL1A1, COL3A1, MATN1*), and growth factor (*ITGB1, EGFR, BMPR2, VEGF*) interactions (fig. 4B) with the GO terms positive regulation of cell migration and adhesion, tissue morphogenesis, epithelial cell proliferation, skeletal system development and leukocyte migration (Fig. 4C). These results imply that through releasing ECM to generate cell-to-cell interaction, cartilage promotes epithelial cell proliferation, migration and morphogenesis in AEC which contributes to the formation of blastema.

Notably, the interaction level of C2 with other clusters had a significant increase in the regeneration process, showing an increasing signal conduction among various cell populations contributing to limb morphogenesis.

Cluster 4 (inflammation cluster) mainly interacted with cluster 2, 3 and 6 and had the strongest connectivity in the early stage (3 dpa) of regeneration. These data further proves the vital role that early inflammation has in releasing cytokines to induce cell differentiation, and triggering the blastema formation in regeneration *(Godwin et al., 2013)*.

Overall, the connectivity map revealed abundant interaction information among different cell clusters which can help provided orientation for future research and targeting in this area.

### The pseudo-temporal ordering of individual cells from all clusters

In order to reconstruct the temporal distribution of all clusters, we used Monocle, an unsupervised algorithm for ordering cells by progressing through differentiation that dramatically increases temporal resolution of expression measurements in a model of axolotl limb regeneration along the pseudo-temporal ordering *(Trapnell et al., 2014)*. Monocle compresses all single-cell transcription datasets in a multi-dimensional space into a two-dimension space for projection. Cells within this two-dimension space (represented by dots) are ordered in “pseudo-time”, with a line through all dots/cells showing a path of the developmental trajectory/timeline. The trajectory of our single-cell RNA-seq datasets displayed a clear large central branch, with few minor branches emanating from it (Fig. 5, A and B). In the pseudo-time trajectory, C5 showed an inchoate state while C3 was at the end of the two early branches (branch 2 and 3), mainly containing C1 cells; branch 1, on the other hand, mainly contained cells from C6 and C2 (Fig. 5A, Fig. S4A). The pseudo-time trajectory was almost consistent with the true time with some cells from 0 dpa (unamputated limb cells) in a primitive state, suggesting that some original stem cells may still exist in unamputated limb tissues (Fig. 5B, Fig. S4B). To investigate the cell alterations in the musculoskeletal system in regeneration, we termed the marker genes, including *COL2A1* (collagen type II alpha 1 chain) for chondrocytes, *ACAN* (Aggrecan) for extracellular matrix in cartilaginous tissue, *COL1A1* (collagen type I alpha 1 chain), *COL1A2* (collagen type I alpha 2 chain) for connective tissues including bone, dermis and tendon, *THBS4* for tendon cells and *POSTN* (Periostin) for osteoblasts or periosteum (Fig. 5C; fig. S4C). Results showed that them all slightly fluctuated in the early period and then increased remarkably during the middle and later periods. To observe more detailed changes in musculoskeletal development-associated signaling factors during regeneration, we investigated the expressional changes of the BMP, FGF and, TGF-β signaling pathway members along the pseudo-time (Fig. 5C, Fig. S4C); results showed that members of FGF and BMP families were expressed with a rising trend. In contrasts, TGF-β signaling pathway members *FOSB* and *ZFYVE16* decreased at the early stage then re-increased at the later stage, *JUNB* was stably expressed with small fluctuations, while other members continuously increased during regeneration pseudo-time. These findings can provided a reference for future regeneration signaling pathway research. We also inspected some specific blastemal-related genes *PRRX1 (Satoh et al., 2011)*, *CIRBP*, and *KAZALD (Bryant et al., 2017)*. *PRRX1* was highly expressed during middle and late stage, while *CIRBP* gradually increased from the early to late stage, and *KAZALD* showed a high level of expression in both the early and late stages (Fig. 5C, Fig. S4C).

**Figure 5.**
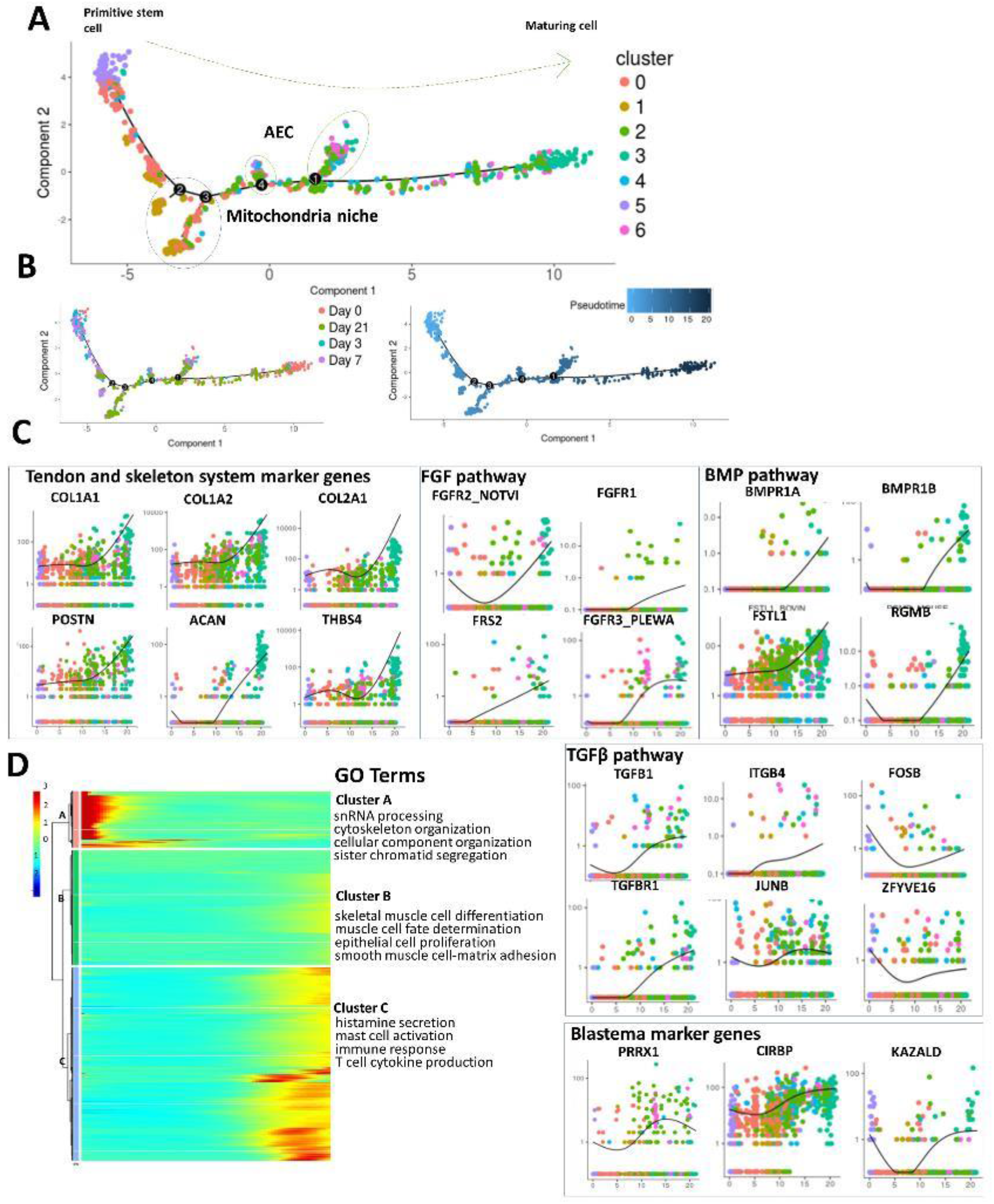
Pseudo-temporal ordering of individual cells. (A) Pseudotime ordering of single cells using 2D PCA. Each data point represents a single cell colored by figure3a clusters age collected. (B) Pseudotime ordering of single cells using 2D PCA. Each data point represents a single cell colored by post amputation time point collected. (C) Expression profiles of Tendon and skeleton system marker genes, different cell signaling pathway components and blastema marker genes along pseudotime ordered by 7 clusters. (D) Heatmap shows the gene expression dynamics during limb regeneration (left). Genes (row) are clustered and cells (column) are ordered according to the pseudotime development. Gene clusters A, B and C were selected for GO analysis (right).

To determine cellular states within regeneration, we further performed clustering analysis. Here, we placed all single cells in their pseudo-time order by compressing small branches into a larger central branch and performed k-means clustering on all genes (p-value <0.05) (Fig. 5D). According to the expressing trend of genes, this approach revealed three gene clusters, hereafter labeled cluster A to C (Fig. 5D); to better understand the biological significance of these three gene clusters, we performed GO analysis (Fig. 5D, right). Cluster A was highly expressed in the early stage with the terms related to cellular component organization and sister chromatid segregation, and both were associated with cell proliferation. Cluster B was expressed during the late stage with GO terms including mature tissue cells differentiation and determination. Lastly, the middle to late stage genes were occupied by cluster C, involving gene expressions relating to immune response. Overall, these findings revealed a large number of factors and pathways that are involved in the regulation of axolotl limb regeneration and hopefully direct future functional studies.

### A continuous transcriptional transition of the musculoskeletal system population in response to the stimuli

Our atlas showed that the *COL2A1+* cells were included not only in C3 but also in C2 (with a lower expression compared to C3) (Fig. S2 E), which indicates that the *COL2A1+* cells with different phenotypes persisted throughout the whole regeneration process. To determine the variances of the *COL2A1+* cells and other musculoskeletal system cells in different regeneration stage, we performed PCA on each time-point cells visualized by t-SNE (Fig. 6A). We attempted to identify the marker genes, *COL2A1*, *ACAN*, *COL1A1*, *COL1A2*, *THBS4* and *POSTN*. Cells with mRNAs for these genes were illustrated according to levels of expression using a four color gradient; grey indicates the lowest detectable mRNA to red which indicates the highest detectable mRNA (Fig. 6B to E; Fig. S6D). The chondrocytes population were found to have a high expression of *COL2A1* and *COL1A1/2* (including *COL1A1* and *COL1A2*) at 0 dpa (unamputated limbs) (Fig. 6B). However, *COL1A1/2*+ and *COL2A1*+ cells had different expression patterns during the regeneration process as reflected by tSNE plots. *COL1A1/2*+ cells were significantly expressed in high proportion (*COL1A1* 58%, *COL1A2* 72%) at 0 dpa, then the number decreased at 3 dpa (*COL1A1* 20%, *COL1A2* 21%), and re-increased (*COL1A1* 47%, *COL1A2* 58%) at 7 dpa; the fluctuation then stopped and the *COL1A1/2*+ cell number was maintained (*COL1A1* 55%, *COL1A2* 70%) at 21 dpa (Fig. 6 B to E; fig. S5, B and C). By contrast, the proportion of *COL2A1+* cells was high (72%) at 0 dpa, and then sharply declined (9%) at 3dpa and (2%) at 7dpa, and increased again at (43%) 21 dpa (Fig. 6, B-E; Fig. S5 B, C). Immunostaining analysis also showed that COL1 had higher expression level at 21 dpa in skeleton, muscle, skin, and blastema (fig. S6), further supporting a mechanism whereby dedifferentiation of the cells and COL1 expression may be important for the regeneration.

**Figure 6.**
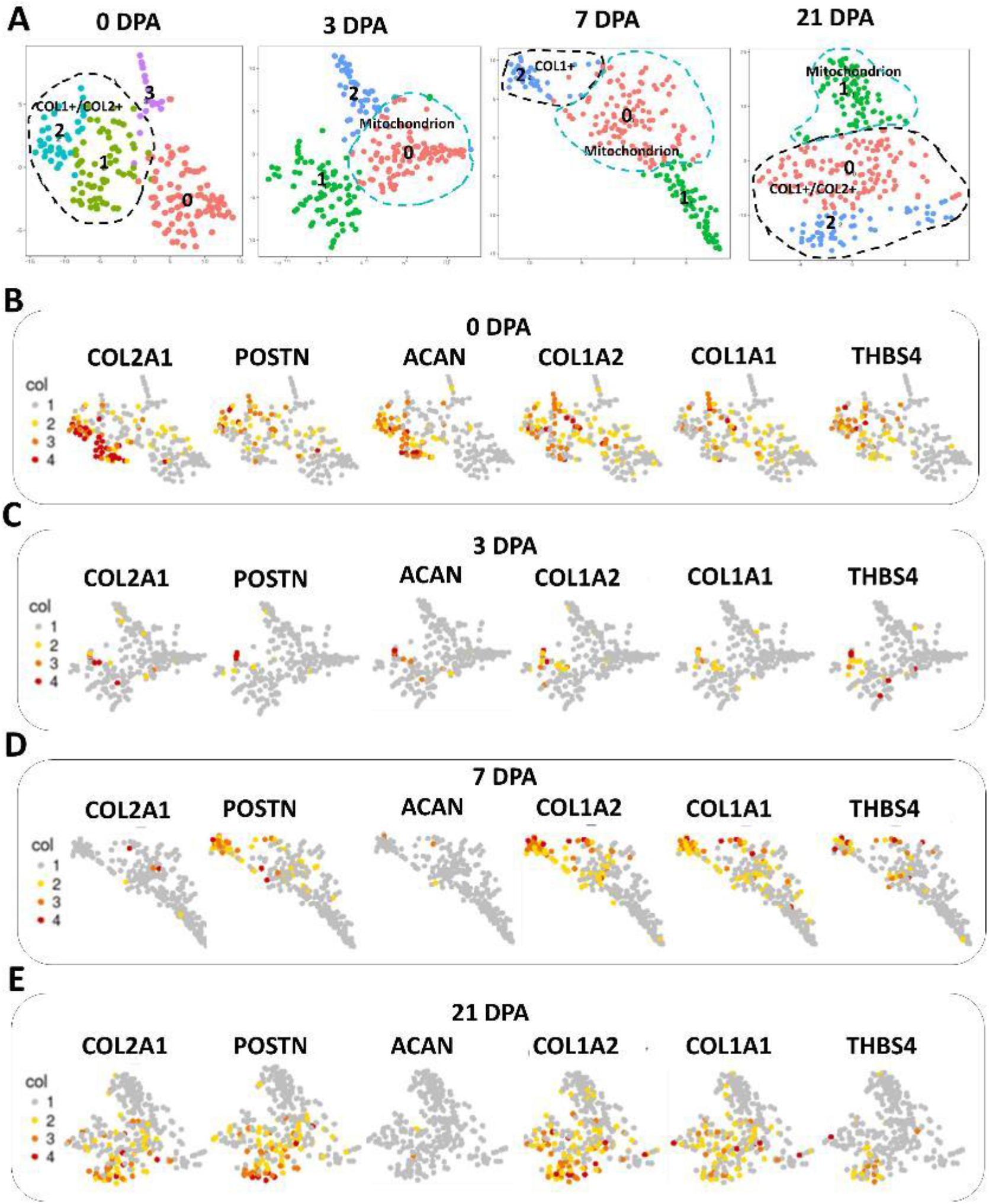
Identification of different lineage cells within 0dpa–21dpa axolotl limbs singe-cell RNA-seq data. (A) t-SNE visualizations of cells at each post-amputation stage. Colors and associated numbers represent individual clusters. (B–E) t-SNE visualization of 0 dpa (B), 3 dpa (C), 7 dpa (D), and 21 dpa. (E) scRNA-seq data overlaid with expression of the cell-type-specific markers COL2A1, COL1A2, COL1A1, THBS4, POSTN and ACAN. Cells are color-coded according to expression level, ranging from low detected to the highest detected levels (grey, yellow, orange and red). Cluster numbers are as shown in (A).

Previous studies have demonstrated that chondrocyte de-differentiation is characterized by the upregulation of type I collagen and downregulation of type II collagen, changing chondrocytes into fibroblast-like cells in vitro *(Cheng et al., 2012)*. In our results, the 0 dpa *COL1A1/2+* and *COL2A1+* cell population had decreased *COL2A1* expression level at 3 dpa showing a rapid reaction to the injury signal. In 7 dpa, the proportion and expression level of *COL1A1/2* re-increased and stabilized at 21dpa while *COL2A1* continued to decline at 7 dpa and rose at 21 dpa. These single-cell level results were consistent with previous studies that the expression of the *COL1A1/2* is earlier than *COL2A1* during limb regeneration *(Asahina et al., 1999; Mitogawa et al., 2015)* and the chondrocytes would dedifferentiated during digits regeneration *(Tanaka et al., 2016)*. This process indicates the chondrocyte dedifferentiation process at the injured site. However, the origins of the re-differentiated *COL2A1+/COL1A1/2+* cells are not clear.

Meanwhile, the proportion of *POSTN+* cells (44% at 0 dpa, 10% at 3 dpa, 22% at 7 dpa, 57% at 21 dpa) and *THBS4+* cells (46% at 0 dpa, 13% at 3 dpa, 12% at 7 dpa, 23% at 21 dpa) were consistent with the expression trend of *COL1A1/2+* cells (Fig. 6, B to E, fig. S5, B and C), indicating that the musculoskeletal cells may regenerate from one cell source.

Conversely, *ACAN*, an integral part of the extracellular matrix in cartilaginous tissue, showed a different expression profile in that it was highly expressed at 0 dpa (34%) then gradually decreased during the regeneration process until 21 dpa (4% at 3 dpa, 1% at 7 dpa, 1% at 21 dpa) (Fig. 6, B-E; Fig. S5 B, C). Further studies are needed to examine the reduction of *ACAN* expression.

### A novel sub-cluster in COL2+ cells

Given these results, we then extracted the COL2+ cells and divided them into 6 sub-clusters with unsupervised seruat clustering methods (Fig. 7A). The sub-cluster 0 and 1 both highly expressed mitochondrion genes, such as ATP6 and COX1 (Fig. 7 B, C). Notably, the sub-cluster 1 specifically expressed mitochondrion genes and was fully contained in the mitochondrion cluster in Fig.2. The sub-clusters 2, 3 and 5 were connective tissue (CT) lineage cells with highly expressed COL1A1. Distinctly, the sub-cluster 3 was labeled mature CT with the marker gene FBN1 (Fig. 7 B-D), while the sub-clusters 2 and 5 were labeled with the marker genes PPRX1 and FBN2 (Fig. 7 B-D), and represented the CT precursors. Meanwhile, in the COL2-CT precursors subpopulation, the sub-cluster 5 highly expressed axolotl upper limb development key gene HOXA13 and HAND2 (Fig. 7 B, C) and showed a strong cell cycling process (Fig S7 B-D), which represented an actively proliferating COL2+ CT regenerative key sub-cluster.

**Figure 7.**
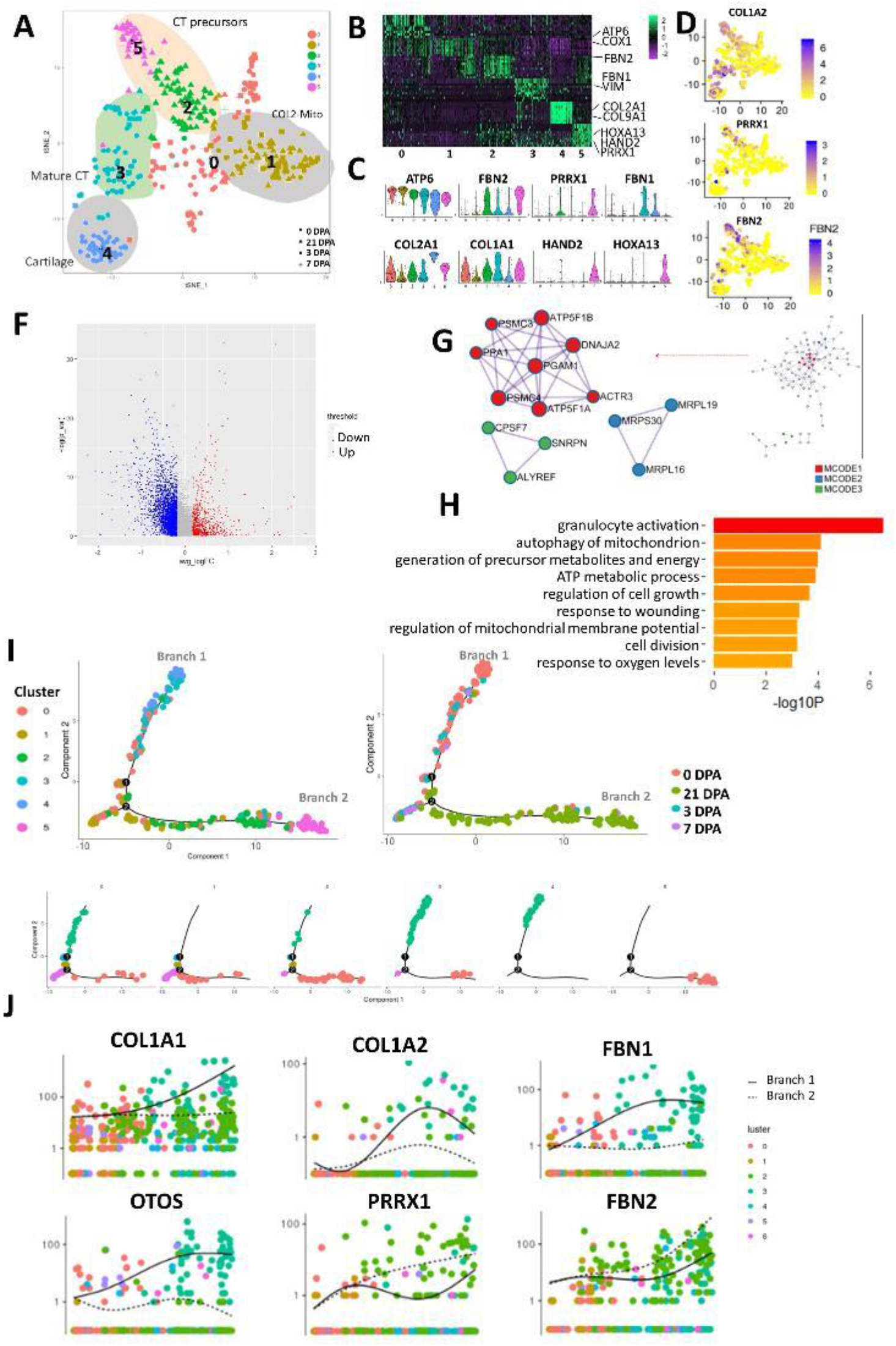
ScRNA-seq analysis reveals COL2-mito sub-cluster in COL2A1+ cells during regeneration process. (A) T-distributed stochastic neighbor embedding (t-SNE) visualizations of COL2A1+ cells sub-clusters identified using the computational pipeline. (B) Heatmap of top 10 differentiated gene of 6 sub-clusters. (C) Violin plots of sub-clusters’ marker genes. (D) Feature Plot of CT marker genes’ expression in individual cells.

As the mitochondria cluster showed the regenerative specificity, we then examined whether the COL2-mito sub-cluster had a regenerative supporting function for the musculoskeletal system. Differential expression analysis was performed between the COL2-mito sub-cluster and the other sub-clusters. The protein-protein interaction enrichment analysis was carried out on the upregulated genes with the databases BioGrid, InWeb_IM, and OmniPath. The molecular complex detection (MCODE) algorithm was applied to identify densely connected network components in the protein-protein interaction network (Fig 7G). In the MCODE network, mitochondrial ATP synthase subunits proteins encoded genes (ATP5F1A, ATP5F1B, PSMC3, PSMC4 et al.) were strongly connected with each other, indicating for an amount of ATP synthases in the COL2-mito sub-cluster. Meanwhile, the mitochondrial ribosomal proteins encoded genes (MRPL16/19/30) and mRNA splicing pathway-associated genes (CPSF7, SNRPN, and ALYREF) were separated connected together, which demonstrated increased mitochondria and a high translation activity. GO analysis indicated that this sub-cluster performed the energy metabolite associated functions such as response to oxygen levels, ATP metabolic process and generation of precursor metabolites and energy (Fig. 7H). Moreover, this sub-cluster showed a response to wounding, such as granulocyte activation and cell division. The pseudo-temporal trajectory of COL2+ cells showed that the mitochondria sub-cluster was located at the started site (fig. S7A) and bifurcated into two big branches; The CT precursors sub-clusters (sub-cluster 2 and 5) were mainly located in branch 1, and the mature CT sub-clusters and cartilage sub-clusters (sub-cluster 3 and 4) mainly composed branch 2 (Fig. 7I). In this trajectory, COL1A1, COL1A2, FBN1 and OTOS were only increasingly expressed in branch 1, while FBN2 expression was higher in branch 2 than branch 1.

## DISCUSSION

In this study, we applied single-cell RNA-sequencing analysis and provided a single-cell transcriptomics which examined the gene expression level of individual cells in axolotl limbs from the early to late stages of regeneration. We computationally identified seven cell clusters in unamputated and regenerated limbs including the first reported mitochondria cluster supporting the regeneration process and a chondrocyte cluster contributing to blastema forming by ECM. We defined the dedifferentiation and re-appearing process of *COL1A1/2+* and *COL2A1+* populations after amputation, and describe the first evidence of how the COL2-mito sub-cluster supported the musculoskeletal system regeneration. by providing energy. These findings provide a single-cell, high-resolution map of the complex cell atlas of adult axolotl limb regeneration.

These data provide several avenues for new research. First, whether chondrocytes contribute to the blastema is yet to be fully resolved. Previous researches of cell lineage tracing did not observe described chondrocytes differentiating in the blastema after amputation. However, our results indicate the chondrocyte cluster showed a strong ligand-receptor interaction with the AEC clusters in all stages which are responsible for cell migration, axon guidance and skeletal system morphogenesis through the connectivity map analysis, which is also consistent with the guidance function of AEC. These results indicating the importance and contribution of chondrocytes in the blastema formation and limb regeneration by ECM, which provide new insights into the connection between cartilage and limb regeneration.

Secondly, the importance of COL2 in regeneration is now more evident. COL2 has been proven not only to be expressed in chondrocytes but has also been detected in tendon-derived stem cells, and synovial-derived stem cells, and has been shown to impact the TGF-b signaling pathway *(Kubosch et al., 2017; Xia et al., 2017; Liu et al., 2018)*. Thus, COL2+ cells present either chondrocytes or multiple lineage stem cells.

Meanwhile, the literature on the dynamics in axolotl connective tissue subtypes reports that dermis bias contributes to the skeleton or dermis *(Gerber et al., 2018)*. In our results, COL1+/COL2+ cells (0 dpa) dedifferentiated into dissimilar cell types with diverse dynamic patterns, indicating COL2+ cells play an essential role in musculoskeletal system regeneration. At 21dpa, COL2A1 reappeared but without the presence of ACAN, which is proof for the earlier expression of COL2 over ACAN, and that the cartilage matrix was not fully formed yet. Our data provide novel evidence for the dedifferentiation and re-differentiation process of the musculoskeletal system from a single-cell level, and may provide valuable clues in the understanding of musculoskeletal system regeneration. Consequently, further research into COL2+ cells’ roll in regeneration and its relationship with other tissues in musculoskeletal system is justified.

In addition to the COL2 functions, several mammalian adult regeneration models have also been studied *(Fernando et al., 2011; Kierdorf and Kierdorf, 2012)*. In axolotl regeneration, the immune responses to amputation and some signaling pathways were similar to the mice digit tips model *(Takeo et al., 2013)*. In contrast, the axolotl rely on the cartilaginous skeleton for regeneration rather than direct ossification as in mice digit tip regeneration *(Han et al., 2008)*. Additionally, the importance of mitochondria in stem cells and liver regeneration in mammals *(Mastrodonato et al., 2012; Khacho et al., 2016)* is similar to the function of our mitochondria cluster in axolotl limb regeneration—an until now undiscovered parallel which may offer further elucidation into energy metabolism in regeneration.

Indeed, previous studies on mitochondria have revealed the links between energy metabolism and transcription rate *(das et al., 2010)*, in addition to clarifying the role of the mitochondria in as well as their roles on the modulation of stem cell differentiation, niche adhesion, and teratoma formation *(Schieke et al., 2008; Wang et al., 2012)*. Tumor cells often contain a distinct mitochondrial profile subpopulation termed cancer stem cells *(Chen and Chan, 2017) (Xie et al., 2015)*. In regeneration, some types of cells act as niche support for stem cells such as nerves *(Kumar and Brockes, 2012)*. In our results, the mitochondria cluster as a specific regenerative cell cluster demonstrated a high translation rate and stem cell functions. This indicates that axolotl limb regeneration relies on a particular cell cluster with a quantity of mitochondria to offer enough energy and respiratory chain intermediates for stress reaction, cell differentiation, and tissue remodeling. Chondrocyte dedifferentiation accompanied by the changes of various genes expressions is known to respond to aspects of the microenvironment, such as oxidative stress *(Ashraf et al., 2016)*. These data suggest that axolotl limbs can could generate a population cells with large amount of mitochondria after amputation, not only deliver energy for a plethora of cellular activities but also to mitigate the oxidative stress which arises in successfully regenerated limbs. It is possible that these same dynamics may be relevant for the clinical treatment on cartilage dedifferentiation and tissue regeneration.

## Conclusion

Here, we reconstructed the first atlas of single-cell gene expression dynamics participating in adult axolotl limb regeneration with the most detailed cell diversity from a single-cell level and uncovered a novel regenerative specific mitochondria-related cell population with likely energy-supplying functions. Our results provided new knowledge on limb regeneration biology, thus revealing novel insights into the regeneration which occurs in the axolotl and other species.

## MATERIAL AND METHODS

### Axolotl husbandry, surgery, and cell isolation

All animal experiments were performed with the approval of the Zhejiang University Ethics Committee (ZJU11602). We established an axolotl colony (*Ambystoma mexicanum*) from ten axolotls obtained from the Guilin Salamander Breeding Base. For all surgical procedures, the animals were anesthetized with 0.03% ethyl 3-aminobenzoate until they were unresponsive to a tail pinch stimulus. We amputated juvenile axolotl right forelimbs at the upper limb level. The 0 dpa cells were harvested from 1mm of adult upper forelimbs. The 3 dpa and 7 dpa cells were harvested from 1mm proximal to the regenerative tissue distal tissues *(Voss et al., 2015)*. After dissection, tissues were finely chopped in DMEM and digested in dissociation solution(C/T, 0.5ml collagenase 2%+10ml, trypsin) at 37°C then centrifuged at 1600 rpm. The precipitation was re-suspended to obtain a concentration of approximately one million cells per milliliter.

### C1 single-cell mRNA sequencing

Blastema and non-regenerating limb stump cells at 5 ×10^5^ cells/ml were loaded onto two 17–25 μm C1 Single-Cell Auto Prep integrated fluidic circuits (Fluidigm) and cell capture was performed according to the manufacturer’s instructions. Individual capture sites were inspected under a light microscope to confirm the presence of single cells. Empty capture wells and wells containing multiple cells or cell debris were discarded for quality control. A SMARTer Ultra Low RNA kit (Clontech) and an Advantage 2 PCR Kit (Clontech) were used for cDNA generation. An ArrayControl RNA Spots and Spikes kit (with spike numbers 1, 4 and 7) (Ambion) were used to monitor technical variability, and the dilutions used were as recommended by the manufacturer. The concentration of cDNA for each single well was determined by Qubit™ dsDNA HS Assay Kit. Multiplex sequencing libraries were generated using the TruePrepTM DNA Library Prep Kit V2 and the Nextera XT Index Kit (Illumina). Libraries were pooled and subjected to sequencing on an Illumina HiSeq 2000 (Illumina).

### Bulk RNA-seq library construction

We used mRNA Capture Beads (VAHTS mRNA-seq v2 Library Prep Kit for Illumina, Vazyme) to extract mRNA from total RNA. PrimeScript™ Double Strand cDNA Synthesis Kit (TaKaRa) was used to synthesize doublestranded cDNA from purified polyadenylated mRNA templates. We used TruePrep DNA Library Prep Kit V2 for Illumina (TaKaRa) to prepare cDNA libraries for Illumina sequencing.

### Data analysis pipeline

We mainly referred to the transcriptome from Bryant’s research (2017) and used the other 2 transcriptomes and 1 genome as an aid. The clean reads were obtained from the high quality region (10-60 nt) of the sequenced Illumina raw reads, and were mapped to the axolotl transcriptome (609,180 transcripts) through Bowtie. The methods for functional annotation were performed according to the methods published by Bryant (2017) *(Bryant et al., 2017)* and we got 83296 transcripts were ultimately annotated.

### Sequencing data analysis

The differential expression genes (DE genes) were calculated by DEseq2 R package. ScRNA-seq expression data were analyzed with Seurat v2.0.1 (PCA, Cluster, t-SNE and cluster) *(Satija et al., 2015)*. In brief, the Seurat object was generated from digital gene expression matrices. In the standard preprocessing workflow of Seurat, we selected 15608 variable genes for PCA, whereupon we performed cell cluster and t-SNE.

Marker genes of each cell cluster were outputted for GO analysis. GOEAST (http://omicslab.genetics.ac.cn/GOEAST/) was used to perform GO analysis. We used GSEA for GO analysis and signaling pathway analysis. Cell clusters were annotated with the information of marker genes and GO analysis results. Digital gene expression matrices with annotations from Seurat were analyzed by Monocle v2.3.6 (pseudo time analysis). The cell-cell interactions were constructed by R package Circlize 0.4.3 *(Gu et al., 2014)*. The count of cell-cell interactions was based on the ligand-receptor pairings. GSEA was performed using GSEA software. The signaling pathway analysis used the gene sets KEGG, REACTOME and BIOCARTA (http://www.broadinstitute.org/gsea/doc/GSEAUserGuideFrame.html).

### Histological examination

The harvested specimens were immediately fixed in 4% (w/v) paraformaldehyde in phosphate-buffered saline for 24 hours. The samples were then dehydrated through an alcohol gradient, cleared, and embedded in paraffin blocks. Histological sections (8 mm) were prepared using a microtome and subsequently stained with hematoxylin and eosin (H&E), and Safranin O.

### Immunostaining

Paraffin sections for immunohistochemistry were treated with 0.05% Trypsin-EDTA (Gibco), 3% (v/v) hydrogen peroxide in methanol, and 1% (w/v) BSA. After overnight incubation at 4℃ with primary antibodies for TOMM20 (Abcam, ab78547), COL1 (Santa Cruz, sc-8784), sections were then incubated with secondary antibodies (Beyotime Biotechnology, China) for 2 hours at room temperature. The DAB substrate system (ZSGB-bio, Beijing, China) was used for color development. Hematoxylin staining was utilized to reveal the cell nuclei.

### Transmission Electron Microscopy

The samples (tissues from 0 dpa, 3 dpa, 7 dpa, and 21 dpa axolotl limbs) were fixed and handled by standard procedures as described previously *(Chen et al., 2012)*. Mitochondria were counted per 30000× picture and each sample was averaged from 10 pictures and analyzed at least three times. Image-Pro Plus (IPP 6.0, Media Cybernetics, Rockville, MD, http://www.mediacy.com.cn/cn/index/index.asp) software was used to counts the mitochondria numbers.

## AVAILABILITY

The raw sequencing data will be available before the publication of this paper.

## ACKNOWLEDGMENTS

This work was supported by the National key research and development program of China [2017YFA0104902]; and the National Natural Science Foundation of China (NSFC) grants [81522029, 81772418, 31570987, 81330041, 81401781, 81572157]. Fundamental Research Funds for the Central Universities.

