## Supplementary material for "Single-Cell RNA-Seq Reveals Novel Mitochondria-related Musculoskeletal Cell Populations during Adult Axolotl Limb Regeneration Process": fig. S

### Supplement figure
#### Figure 1S

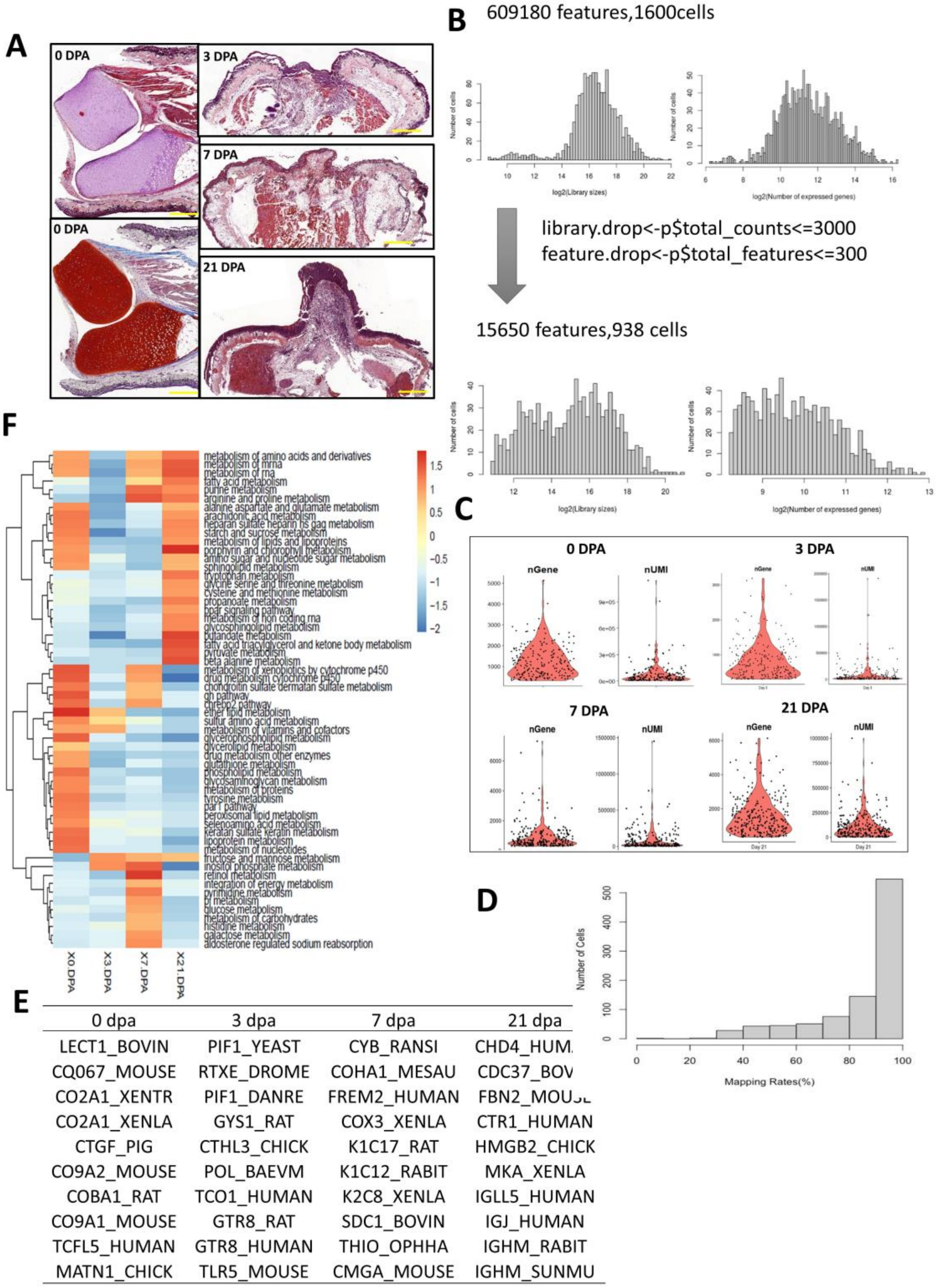


| 0 dpa | 3 dpa | 7 dpa | 21 dpa |
|---|---|---|---|
| LECT1_BOVIN | PIF1_YEAST | CYB_RANSI | CHD4_HUM[illegible] |
| CQ067_MOUSE | RTXE_DROME | COHA1_MESAU | CDC37_BOV[illegible] |
| CO2A1_XENTR | PIF1_DANRE | FREM2_HUMAN | FBN2_MOU[illegible] |
| CO2A1_XENLA | GYS1_RAT | COX3_XENLA | CTR1_HUMAN |
| CTGF_PIG | CTHL3_CHICK | K1C17_RAT | HMGB2_CHICK |
| CO9A2_MOUSE | POL_BAEVM | K1C12_RABIT | MKA_XENLA |
| COBA1_RAT | TCO1_HUMAN | K2C8_XENLA | IGLL5_HUMAN |
| CO9A1_MOUSE | GTR8_RAT | SDC1_BOVIN | IGJ_HUMAN |
| TCFL5_HUMAN | GTR8_HUMAN | THIO_OPHHA | IGHM_RABIT |
| MATN1_CHICK | TLR5_MOUSE | CMGA_MOUSE | IGHM_SUNMU |

**Figure. S1 Single-Cell RNA-Seq data features and changes during axolotl limb regeneration the time course.**

(A) HE staining of samples at 0, 3, 7, and 21dpa. (B) Frequency distribution histogram of the genome mapping rate (C) Violin plots of the number of genes and transcripts detected per cell at each day post-amputation. (D) Top 10 of differentially expressed genes of 4 time points group. (E) Heatmap of metabolism-associated signaling pathway by GSEA. Heatmap of BMP, FGF, TGFb, Wnt and MAPK signaling pathway by GSEA.

#### Supplement figure

#### Figure S2

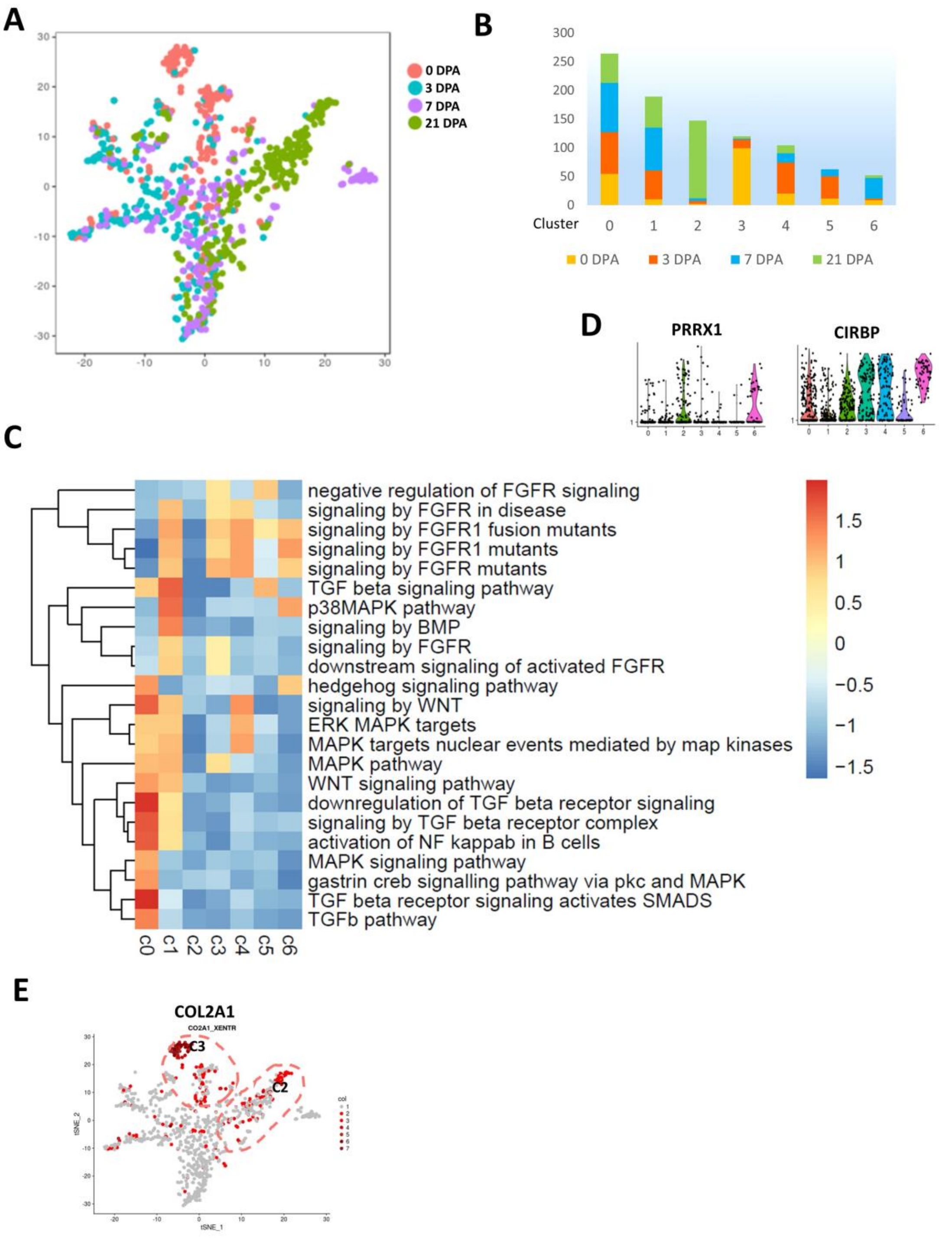


**Figure S2. Cell heterogeneity in axolotl limb regeneration tissues.**

(A) T-distributed stochastic neighbor embedding (t-SNE) visualizations of four days post-amputation identified using the computational pipeline (B) Histogram of the proportion of four days post-amputation cells in 7 clusters. (C) Heatmap of musculoskeletal system associated signaling pathway in 7 clusters by GSEA. (D) The expression of blastemal specific gene in 7 clusters. (E) Feature Plot of COL2A1 expression in total individual cells.

#### Supplement figure

#### Figure S3

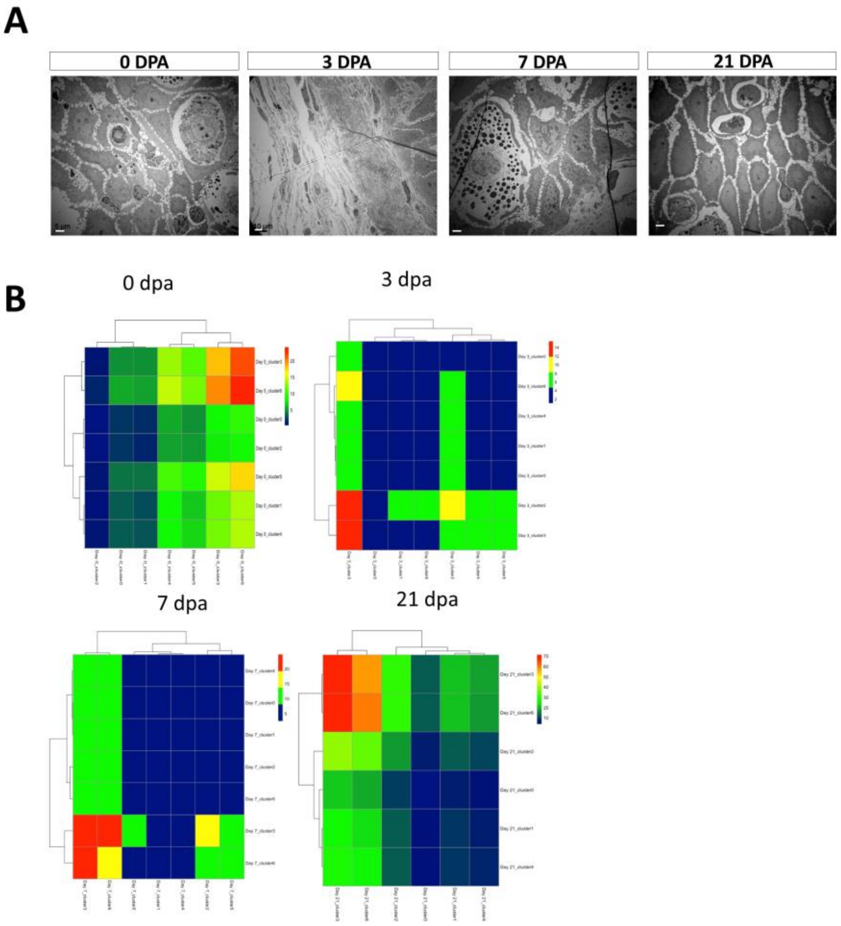


**Figure S3. Connectivity map reveals interactions among clusters.**

(A) TEM of regenerative samples at each time points. Scale bar, 5 μm. (B) Heatmap shows the total numbers of putative receptor-ligand interactions between two sub-clusters from four time points post-amputation.

#### Supplement figure

#### Figure S4

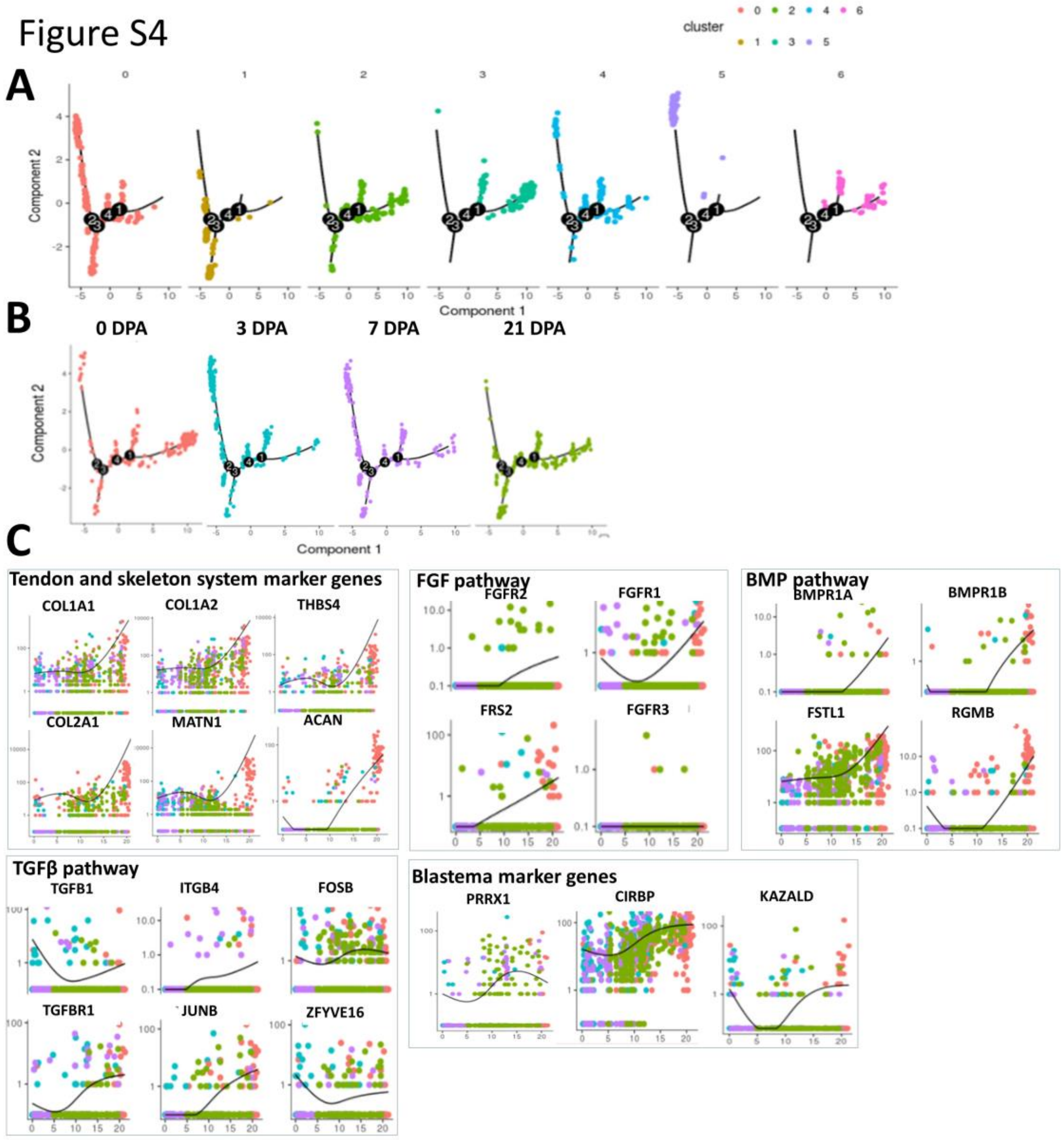


**Figure S4. Pseudo-temporal ordering of cells from all time points.**

(A) Pseudotime ordering of single cells using 2D PCA. Each data point represents a single cell colored by figure3a clusters age collected. (B) Pseudotime ordering of single cells using 2D PCA. Each data point represents a single cell colored by post amputation time point collected. (C)Expression profiles of Tendon and skeleton system marker genes, different cell signaling pathway components and blastema marker genes along pseudotime ordered by 4 days post-amutation.

Supplement figure

Figure S5

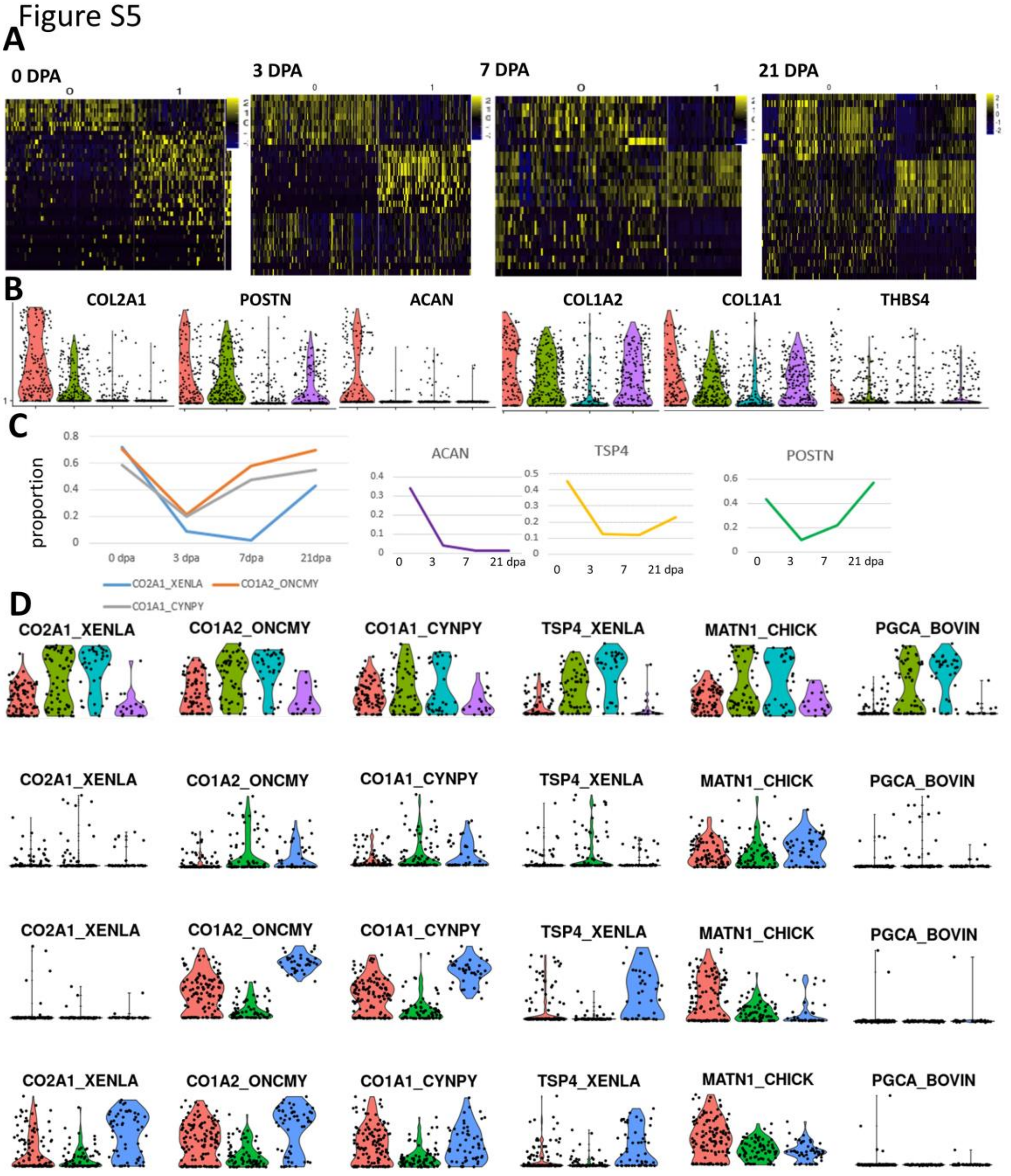


**Figure S5. Identification of different lineage cells within 0dpa–21dpa axolotl limbs Singe-cell RNA-seq Data**

(A) Heatmap of 0 dpa, 3 dpa, 7 dpa, and 21 dpa differentiation expression top10 genes. Heatmap are color-coded according to expression level, ranging from low detected to the highest detected levels (blue and yellow). (B) Violin plots of marker genes COL2A1, COL1A2, COL1A1, THBS4, POSTN and ACAN in each subclusters from 4 post-amputation time points. (C) Line charts showed the proportions of COL2A1+, COL1A2+, COL1A1+, THBS4+, POSTN+ and ACAN+ cells in each day post-amputation. (D-G) Violin plots of marker genes COL2A1, COL1A2, COL1A1, THBS4, POSTN and ACAN in each subclusters respectively from 0dpa, 3dpa, 7dpa and 21dpa. Cluster numbers are as shown in figure2 (A).

Supplement figure

Figure S6

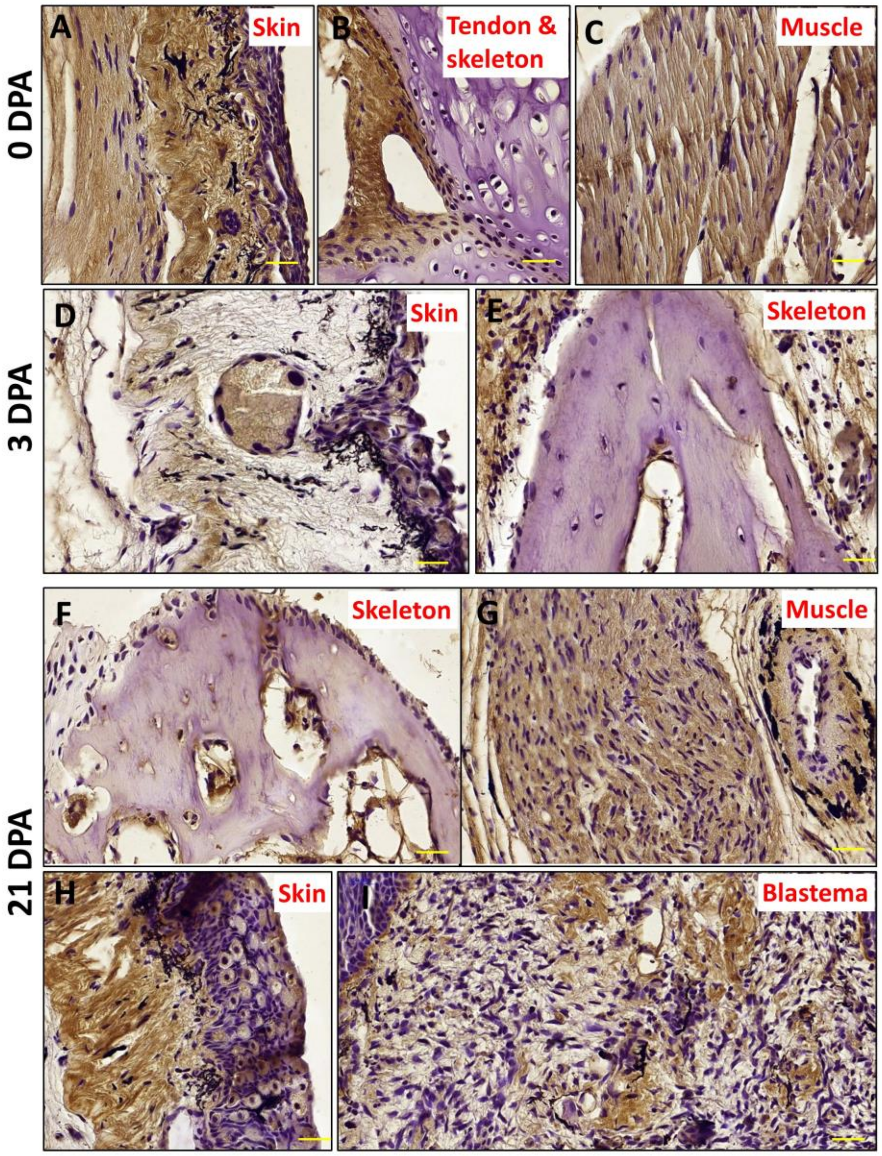


**Figure S6. Validation of COL1 variance in regeneration.**

Immunostaining of COL1 in 3 dpa and 21 dpa axolotl limb distal tissues. COL1 in (A) skin, (B) tendon and skeleton,(C) muscle of 0 dpa tissues, (D)skin and (E) skeleton of 3 dpa tissues, and (F)skeleton, (G) muscle, (H) skin and (I) blastema of 21 dpa tissues.

Supplement figure

Figure S7

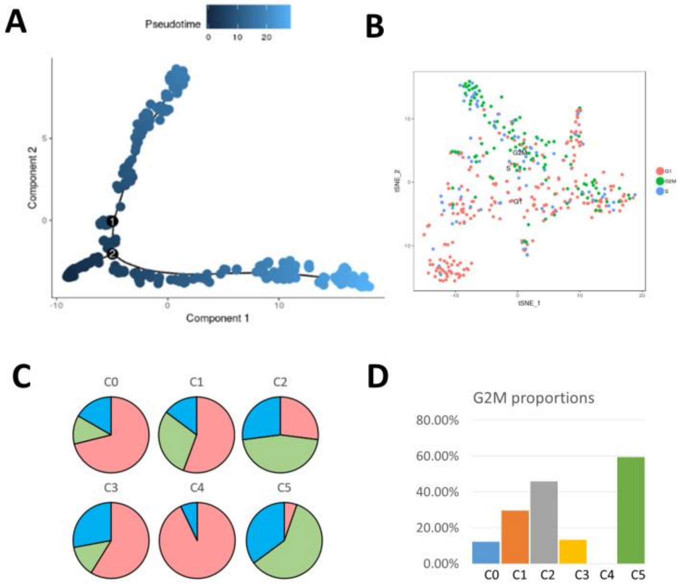


**Figure S7. Proliferation analysis of COL2+ sub-clusters.**

(A) Pseudotime ordering of single cells using 2D PCA. Each data point represents a single cell colored by pseudotime. (B) Proliferation analysis of 6 COL2+ sub-clusters. (C) Proportions of S, G1 and G2/M periods in each sub-clusters proliferation analysis. (D) Proportions of G2/M period in each sub-clusters proliferation analysis.
